# Viral genetic diversity and functional potential in polar and subarctic sea ice

**DOI:** 10.64898/2025.12.12.693969

**Authors:** Kalle J. Pettersson, Tatiana Demina, Eeva Eronen-Rasimus, Simon Roux, Sirja Viitamäki, Igor S. Pessi, Hanna M. Oksanen, Philipp Assmy, Hermanni Kaartokallio, Jenni Hultman

**Author notes:** Corresponding author: Tatiana Demina. Shared first authorship (Kalle J. Pettersson, Tatiana Demina).

## Abstract

Sea ice plays a critical role in regulating the global climate and serves as a unique habitat for diverse microbial communities. Still, our understanding of viruses in these communities remains limited. To further uncover the diversity and functional potential of viruses in polar and subarctic sea ice, we explored the viral component of Arctic, Baltic Sea, and Antarctic sea ice metagenomes. Altogether, 550 viral operational taxonomic units (vOTUs) were recovered, most of which were putatively classified within the class *Caudoviricetes*, which comprises bacterial and archaeal tailed double-stranded DNA viruses. Hosts were predicted for 187 vOTUs, with *Gammaproteobacteria* and *Bacteroidia* being the most prevalent viral host groups. Potential functions were assigned for 56% of predicted viral gene products, including putative auxiliary metabolic genes (AMGs) involved in oxidative metabolism, photosynthesis, and metabolism regulation under stress conditions. Related viral genomes carrying similar AMGs were detected in other Arctic and more geographically distant freshwater, marine, and ice environments. Genus- and/or family-level links between the studied vOTUs were detected across samples. Our results suggest diverse and complex virus-host interactions in sea ice and highlight the essential roles viruses may play in sea ice ecosystem dynamics across polar and subpolar environments.

## Introduction

Sea ice is an important factor regulating the ecology of polar marine ecosystems and global climate (Dieckmann and Hellmer 2010), and its decline in the Arctic Ocean is a key indicator of climate change (Fox-Kemper et al. 2023). The Arctic Ocean has experienced a long-term decrease in sea ice extent, and in the Antarctic, a striking decrease in sea ice extent has been observed in recent years (Parkinson and DiGirolamo 2021; Purich and Doddridge 2023; Abram et al. 2025). The Baltic Sea, with brackish water, is covered annually with a varying extent of sea ice (Granskog et al. 2006; Vihma and Haapala 2009), having an overall decreasing trend in maximum sea ice extent over the past decades (Haapala et al. 2015; Kaartokallio et al. 2025). Predicting the consequences of sea-ice loss on local ecosystems remains difficult (Post et al. 2013; Naakka et al. 2025).

Sea ice is a complex environment comprising a solid ice matrix with a network of small channels and pockets filled with hypersaline brines. Brine channel habitats are tightly packed and biofilm-like, as pores are often filled with extracellular polysaccharide gels where various organisms are embedded, including algae, bacteria, archaea, viruses, fungi, micro- and macrograzers (Arrigo 2017; Caron et al. 2017; Bluhm et al. 2017; Lund-Hansen et al. 2024; Rapp et al. 2025). Sea ice microbial communities initially form through particle scavenging during ice formation, followed by enrichment and community development, where cold-adapted species become dominant (Deming and Collins 2017; Lund-Hansen et al. 2024). Typically, heterotrophic bacteria of the classes *Flavobacteria* and *Gammaproteobacteria*, and to a lesser extent *Alphaproteobacteria, Verrucomicrobia* and *Bacilli*, dominate in bacterial communities in polar sea ice (Boetius et al. 2015). In Baltic Sea ice, bacterial communities are similar to those in polar sea ice (Eronen-Rasimus et al. 2015).

Sea ice brines are enriched with viruses, where they experience higher contact rates with their hosts compared to those in seawater (Wells and Deming 2006; Maranger et al. 1994). So far, sea ice virus isolates have been obtained from polar and Baltic Sea ice on *Flavobacterium, Colwellia, Octadecabacter, Glaciecola*, and *Pseudoalteromonas* strains (Borriss et al. 2003; Yu et al. 2014; Luhtanen et al. 2014, 2018). These are all tailed phages (*Caudoviricetes*), except f327, a filamentous phage from Arctic sea ice (Yu et al. 2014). The known sea ice virus isolates are host-specific (Borriss et al. 2003; Luhtanen et al. 2014; Senčilo et al. 2015; Luhtanen et al. 2018) and their infection results in the lysis of the host cell under laboratory conditions (Senčilo et al. 2015; Demina et al. 2022). Their genome sequences are diverse and unique, not closely related to one another (Borriss et al. 2007; Senčilo et al. 2015; Demina et al. 2022), except that *Shewanella*-infecting phages 1/4 and 1/40 are suggested to belong to the same genus (Senčilo et al. 2015).

Meta-omics approaches targeting viruses have been applied to various polar environments (Heinrichs et al. 2024), but only a few such studies are available specifically for polar sea ice (Zhong et al. 2020, 2023; Liu et al. 2022; Kanaan and Deming 2025). Viral communities in sea ice and underlying seawater appear to overlap, but are distinct from those found in other ecosystems (Zhong et al. 2023; Liu et al. 2022). Based on metatranscriptomics, active viral infections were observed in *Polaribacter* which dominated in subzero Arctic sea ice brines (Zhong et al. 2023). Diverse auxiliary metabolic genes (AMGs) predicted in viral sequences from both Arctic and Antarctic sea ice suggest that viruses may modulate bacterial metabolic processes, including cold adaptation, in these environments (Zhong et al. 2020; Liu et al. 2022).

Altogether, previous studies suggest that sea ice is a hot spot of viral diversity, which remains undersampled. To further uncover the diversity of viruses in polar and subarctic sea ice, we analysed viral sequences in bulk metagenomes originating from Arctic, Baltic Sea, and Antarctic sea ice. Given that sea ice remains an undersampled environment, we hypothesized that viral sequences from these habitats would encode products with largely unknown functions and share limited similarity with previously described sea ice viruses. Consistent with this, predicted viral operational taxonomic units (vOTUs), which mostly represented tailed phages that were putatively linked to abundant sea ice bacteria, showed previously undescribed genetic diversity.

## Materials and Methods

### Sampling and sequencing

A total of nine sea ice samples were used in this study: three were collected in the Arctic Ocean north of Svalbard in early summer 2015 (this study, Assmy et al., 2017), three in the Baltic Sea in winter 2006 (Eronen-Rasimus et al., 2015), and three in the western and eastern Weddell Gyre, Antarctica, in winter 2013 (Table S1, Fig. 1, Eronen-Rasimus et al., 2017).The samples were processed as described in (Eronen-Rasimus et al. 2015; Assmy et al. 2017; Eronen-Rasimus et al. 2017; Luhtanen et al. 2018). One of the samples collected during the Antarctic winter expedition (AWECS, leg ANT-XXIX/6, sample 24) was exceptional since it was the middle part from a 3-m thick ice flow, which had a strong odour of hydrogen sulfide (H_2_S), suggesting anoxic conditions (Eronen-Rasimus et al. 2017). For other Antarctic and Baltic sea ice samples, all collected ice layers were pooled together, while for Arctic samples, only the top and bottom 20 cm were pooled together. Antarctic and Baltic Sea ice samples were melted at +4 °C and meltwater was filtered through 0.2 μm membrane filters (Whatman) immediately after melting during the cruises. For Arctic samples, ice was stored at -20°C during the cruise, shipped to Finland at the beginning of October 2015, melted at +4 °C and filtered. The filters were stored at -80°C until DNA extraction. DNA from the filters was extracted within one month after sample arrival to Finland using PowerSoil DNA Isolation Kit (MoBio Laboratories Inc., Carlsbad, CA, USA) as described in (Eronen-Rasimus et al. 2017). Metagenomic libraries were constructed with NexteraXT kit (Illumina) and sequenced with NextSeq 500 (150 + 150 bp) in the Institute of Biotechnology, University of Helsinki, Finland. The raw metagenomic data for this study have been deposited in the European Nucleotide Archive (ENA) at EMBL-EBI under the accession number PRJEB101482.

**Figure 1.**
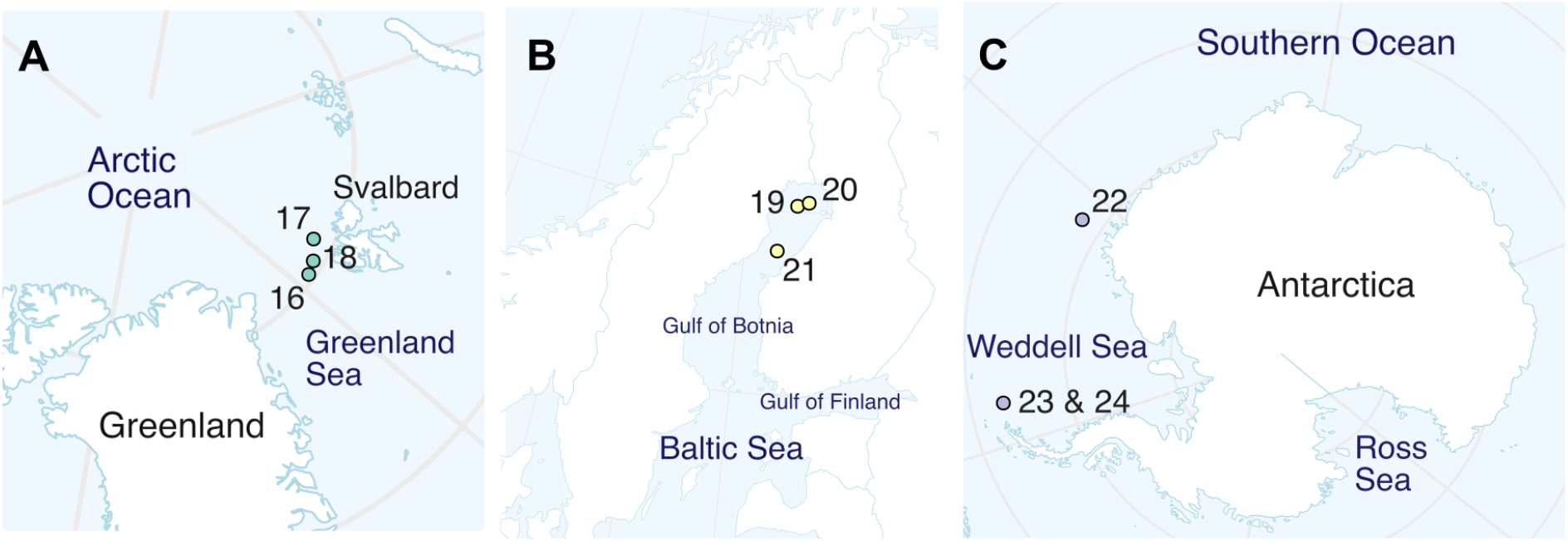
Sampling locations for the (A) Arctic, (B) Baltic Sea, and (C) Antarctic sea ice samples used in this study.

### Taxonomic profiling of bacteria, archaea, and eukaryotes

Metagenomic reads were trimmed using Cutadapt v. 3.5 (-q 30 -m 50) (Martin 2011), and fastp v. 0.23.4 (-g -l 50) was used to trim polyG tails (Chen 2023; Chen et al. 2018). Read quality control was performed with FastQC v. 0.11.9 (Andrews 2010). To study the taxonomy of bacteria and archaea in the samples, read-based analyses of unassembled SSU rRNA gene sequences were performed with phyloFlash v. 3.4.2 (Gruber-Vodicka et al. 2020) using SILVA 138.1 SSURef NR99 database (main taxonomic ranks only) (Seah and Team 2023), followed by Phyloseq v. 1.41.1 (McMurdie and Holmes 2013) analysis in R (https://cran.r-project.org). Hits to mitochondria, chloroplasts, and vertebrates were filtered out. The data were then normalized by dividing the counts by their sum to account for differences in library size between the samples. Principal coordinates analysis (PCoA) was performed to study the differences between host community structures without eukaryotes, using Bray-Curtis dissimilarity index to compute the distances. These differences were then confirmed using permutational ANOVA (PERMANOVA) with 9999 permutations, using R package vegan v. 2.6.4 (https://github.com/vegandevs/vegan, https://cran.r-project.org).

### Assembling metagenomes and identifying viral sequences

MetaSPAdes v. 3.15.5 was used for the assembly of trimmed reads, followed by the quality control with MetaQUAST v. 5.2.0 (Mikheenko et al. 2016; Nurk et al. 2017). Bowtie v. 2.4.4 (Langmead and Salzberg 2012) was used to map the metagenomic reads onto the assemblies. Viral contigs were identified with geNomad v. 1.5.2 (Camargo et al. 2024), and their quality was assessed with CheckV v. 1.0.3 (Nayfach, Camargo et al. 2021). Contigs that were ≥ 5 kb or ≥ 50% complete, had at least one viral gene predicted, and host to viral genes ratio < 1.5 were selected and dereplicated across all samples using the scripts anicalc.py and aniclust.py from CheckV (95% average nucleotide identity and 85% alignment fraction) (Nayfach, Camargo et al. 2021). Since some vOTUs showed similarities to algal retrotransposons in BLASTn searches (core_nt database, E-value threshold of 1e-3), we additionally used TEsorter v. 1.4.7 with the default REXdb database (viridiplantae v.3 plus metazoa v.3) (Zhang et al. 2022) to identify putative transposable elements (TEs). These TEs were then excluded from the final set of vOTUs. All vOTUs, except seven classified as eukaryotic viruses (*Megaviricetes*), were annotated with Phold v. 0.2.0 (Bouras et al. 2026), using Foldseek v. 9.427df8a (van Kempen et al. 2024), ProstT5 (Heinzinger et al. 2024), as well as Colabfold v. 1.5 (Mirdita et al. 2022) as core dependencies, and the PHROG database (Terzian et al. 2021). Genes assigned to the PHROG category “moron, auxiliary metabolic gene and host takeover” in the Phold annotations were assumed to be potential AMGs, some of which were explored in more detail by searching the vOTUs of interest against the IMG/VR v.4 database (Camargo et al. 2023) and/or NCBI core_nt database using BLASTn (E-value threshold of 1e-5) (Altschul et al. 1990). Hits used for further analyses were additionally annotated with Phold v. 0.2.0 (Bouras et al. 2026). Pairwise viral genome comparisons were visualized with Easyfig v. 2.2.2, where BLASTn E-value threshold of 1e-3 was used (Sullivan et al. 2011). Lifestyle for all vOTUs except *Megaviricetes* was predicted with BACPHLIP v. 0.9.6 using the 95% confidence threshold (Hockenberry and Wilke 2021). iPHoP v. 1.3.3 with the Aug_2023_pub_rw database was used to predict bacterial and archaeal hosts for vOTUs (Roux et al. 2023). The score thresholds of 90 and 75 were applied for the genus- and family-level predictions, respectively. In cases of a single vOTU having multiple host predictions, the prediction with the highest confidence score was preferred.

### Viral communities

Metagenomic reads were mapped to vOTUs with Bowtie v. 2.4.4 (Langmead and Salzberg 2012), sorted and indexed with SAMtools v. 1.16.1 (Danecek et al. 2021), and the number of reads mapping to each vOTU (≥95% identity and ≥75% coverage) as well as horizontal coverage were counted with CoverM v. 0.6.1 (https://github.com/wwood/CoverM). Abundance values normalized to reads mapped per kilobase of contig per million reads (RPKM) were visualized as heatmaps using ggplot2 v. 3.4.4 in R (https://ggplot2.tidyverse.org/reference/ggplot2-package.html, https://cran.r-project.org). Clustering vOTUs into family- and genus-level clusters based on AAI was done as described in https://github.com/snayfach/MGV/tree/master/aai_cluster (Nayfach, Paez-Espino et al. 2021). Statistical analyses were done with vegan v. 2.6.4 in R (https://github.com/vegandevs/vegan, https://cran.r-project.org). Differences in vOTU abundances across samples were analysed using PCoA and confirmed with permutational ANOVA (PERMANOVA) with 9999 permutations. Distances were computed using the binary Jaccard dissimilarity metric. In all analyses involving RPKM values, a relaxed horizontal coverage threshold of 10% was used to maximize the capturing of low-abundance viruses (Trubl et al. 2025).

### Comparisons of virus isolates and metagenomic data

Metagenomic reads were mapped against 11 complete genome sequences of sea ice virus isolates (Table S2) using Bowtie v. 2.4.4 (Langmead and Salzberg 2012). Isolate sequences were also compared to metagenomic contigs using BLASTn (whole genomes and ORFs) or BLASTp (ORFs’ amino acid translations) (Altschul et al. 1990; Camacho et al. 2009). For that, Prodigal v. 2.6.3 was used to predict ORFs and their amino acid translations in contigs (Hyatt et al. 2010). Sequence similarities were visualized using the Circoletto web tool with the BLASTn E-value threshold of 1e-2 (Darzentas 2010).

### Whole genome network-based comparisons with vConTACT2

The vOTUs with a length of ≥ 10 kbp were compared against different datasets using whole-genome gene-sharing profiles with vConTACT2 v. 0.9.19 and the NCBI Bacterial and Archaeal Viral RefSeq V201 database (Bin Jang et al. 2019). In addition, viral contigs from Arctic sea ice and cryopegs (Zhong et al. 2020) and Antarctic under-ice waters (Lopez-Simon et al. 2023), which were selected based on their length (≥ 10 kbp) and dereplicated in the same way as the vOTUs from this study, were added into the vConTACT2 analysis. Furthermore, the analysis also included 143 high-quality ice-associated uncultivated viral genomes (UViGs) retrieved from the IMG/VR v.4 database with the following search criteria: ≥ 10 kb long, high quality, high confidence, and having “sea ice” as ecosystem or “sea ice”, “ice”, “sea ice brine” as habitat. Cytoscape v. 3.10.0 was used to visualize the output network (Shannon et al. 2003).

## Results and Discussion

### Microeukaryotes, *Pseudomonadota* and *Bacteroidota* dominated across samples

Sequencing of the nine sea ice samples resulted in 14.4-21.8 M raw reads per sample, which were filtered to 10.8-17.4 M reads per sample after trimming (Table S3). Read-based analyses showed that the most abundant bacterial phyla across the samples were *Pseudomonadota* (also known as *Proteobacteria*) and *Bacteroidota* (Fig. 2A). At the class level, *Gammaproteobacteria* contributed to the abundance of the phylum *Pseudomonadota* more than *Alphaproteobacteria*. The phyla *Bacillota* and *Actinomycetota* were prevalent in the Arctic and Baltic samples. The phylum *Planctomycetota* was abundant in the Baltic samples, consistent with previous observations (Eronen-Rasimus et al. 2015). The phyla *Desulfobacterota* and *Campylobacterota* were detected in the Antarctic sample 24, and these phyla are known to include members involved in sulfur metabolism (Waite et al. 2020, 2017, 2018), reinforcing the previous observation of sulfate-reducing bacteria in the same presumably anoxic sea ice sample (Eronen-Rasimus et al. 2017).

**Figure 2.**
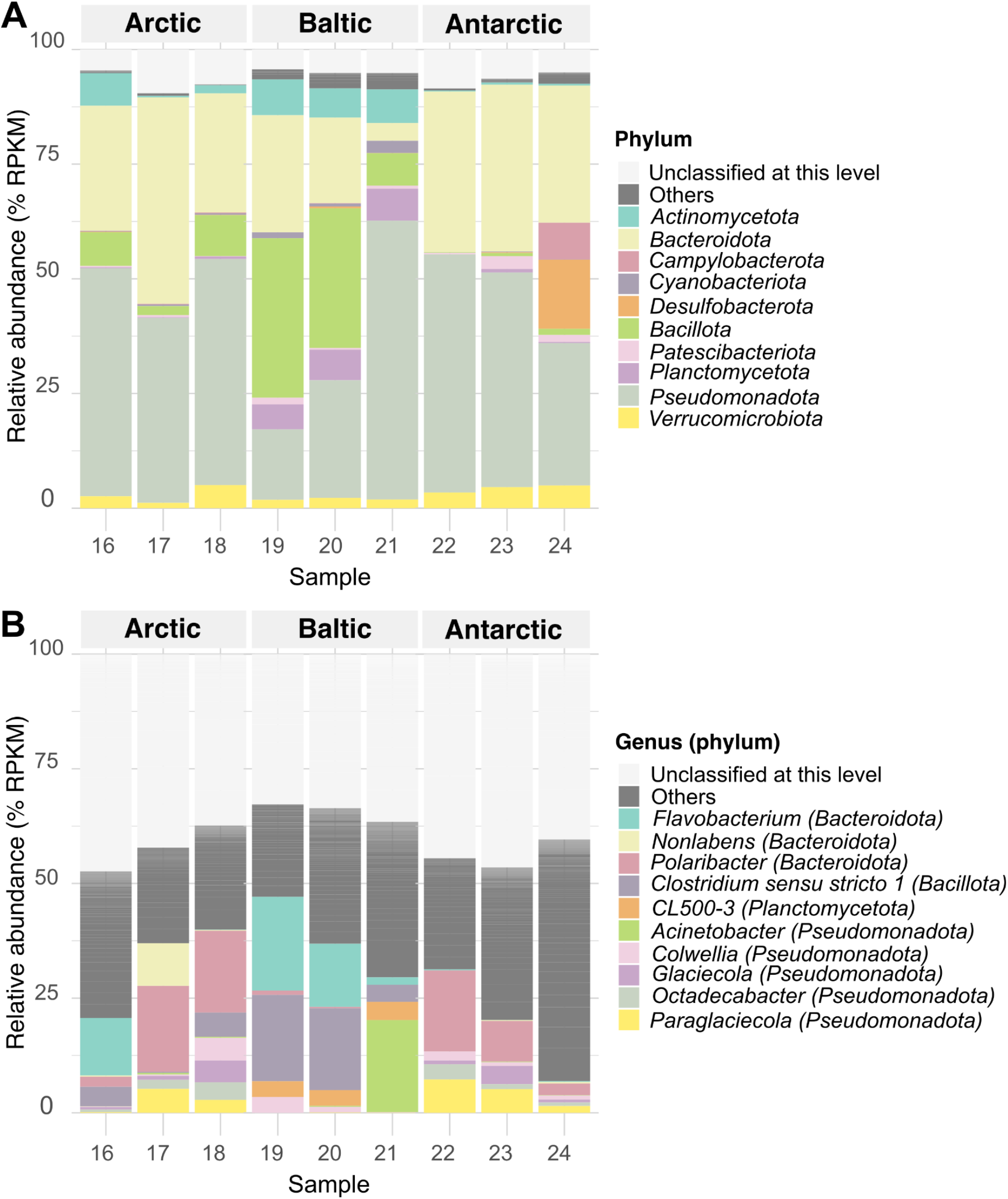
The ten most abundant bacterial and archaeal (A) phyla and (B) genera across samples (all eukaryotes are filtered out). The data were normalized after removing eukaryotic sequences (see Materials and Methods). RPKM, reads per kilobase million

The genera *Polaribacter, Octadecabacter*, and *Flavobacterium* are usually among the most commonly found bacterial groups in sea ice (Rapp et al. 2025). Consistently, the genus *Polaribacter* was highly abundant in the Arctic and Antarctic samples (Fig. 2B). In addition, *Paraglaciecola* and *Octadecabacter* were also abundant in most of the Arctic and Antarctic samples. The genera *Flavobacterium* and *Clostridium sensu stricto 1* were abundant in some of the Arctic and Baltic Sea samples. In addition, *Nonlabens* and *Acinetobacter* had high relative abundance in samples 17 (Arctic) and 21 (Baltic Sea), respectively. From the observed genera, *Clostridium sensu stricto 1* is anaerobic (or aerotolerant) and not found in sea ice, but it is dominant in fecal microbial communities of, e.g., marine seal (Godino et al. 2024), fish (Skrodenytė-Arbačiauskienė et al. 2022), and seabirds (Laviad-Shitrit et al. 2019). Thus, *Clostridium sensu stricto 1* could possibly be transferred to the sea ice from animal feces, likely not representing actual sea ice bacteria.

Compared to the ∼900 different bacterial genera identified, only 14 archaeal ones were detected, which is in line with the observation that archaeal contributions to sea ice total microbial communities are typically small (Deming and Collins 2017). Among unicellular eukaryotes, the most common groups across samples were *Alveolata* and/or *Stramenopiles* (Fig. S1A). The two groups were composed mainly of dinoflagellates and diatoms, respectively (Fig. S1B). Among other eukaryotes, protists (*Rhizaria*), flatworms (*Platyhelminthes*), crustaceans (*Malacostraca* and *Maxillopoda*), as well as fungi were also observed, highlighting a wide diversity of organisms inhabiting sea ice.

In the PCoA analyses, the samples from the Arctic and Antarctica tended to cluster more closely with each other than with those from the Baltic Sea, although the limited number of samples and divergent nature of the datasets precludes robust statistical validation of this pattern (Fig. S2). The samples 17 (Arctic) and 22 (Antarctic) clustered closely with each other. These two samples were characterised by high relative abundance of dinoflagellates and a low relative abundance of diatoms. Interestingly, these two stations were subjected to flooding (Fernández-Méndez et al. 2018; Tison et al. 2017), which is known to replenish nutrients in sea ice (Henley et al. 2023). Typically, diatoms dominate in polar sea ice communities, however, dinoflagellate dominance over diatoms has also been observed, which is predicted to occur due to climate change, i.e., elevated nutrient concentrations, high irradiance and warmer ice temperatures (Lund-Hansen et al. 2024).

### Most vOTUs were predicted to be tailed viruses from the class *Caudoviricetes*

Assembling of the metagenomes resulted in 29.3-198.3 K contigs per sample (Table S3). Contigs predicted as viral by geNomad were further filtered based on their size, completeness, and the number of viral genes (see Materials and Methods). From 624 dereplicated presumably viral contigs, 74 were predicted as other TEs, mostly LTR Gypsy retrotransposons, by TEsorter (Table S4). These TE contigs were removed, and the final set of vOTUs included 550 contigs, with each vOTU consisting of a single contig (Table S5). The vOTUs were 3-71 kbp long, with the median length of ∼7 kbp. Five vOTUs were 99-100% complete, including one putative provirus.

Most vOTUs (537) were assigned to a taxonomic unit by geNomad (Table S5, Fig. 3A). From the classified vOTUs, the majority (528) were placed within the class *Caudoviricetes*, which comprises tailed dsDNA bacterial and archaeal viruses. However, most of these *Caudoviricetes* vOTUs (500) could not be classified further. Still, some vOTUs within *Caudoviricetes* could be putatively classified at the family level: *Schitoviridae* (11), *Kyanoviridae* (4), *Autographiviridae* (3), *Zobellviridae* (2), and *Chaseviridae* (1). In addition to *Caudoviricetes*, the classes *Megaviricetes* (7) and *Malgrandaviricetes* (2) were also observed.

**Figure 3.**
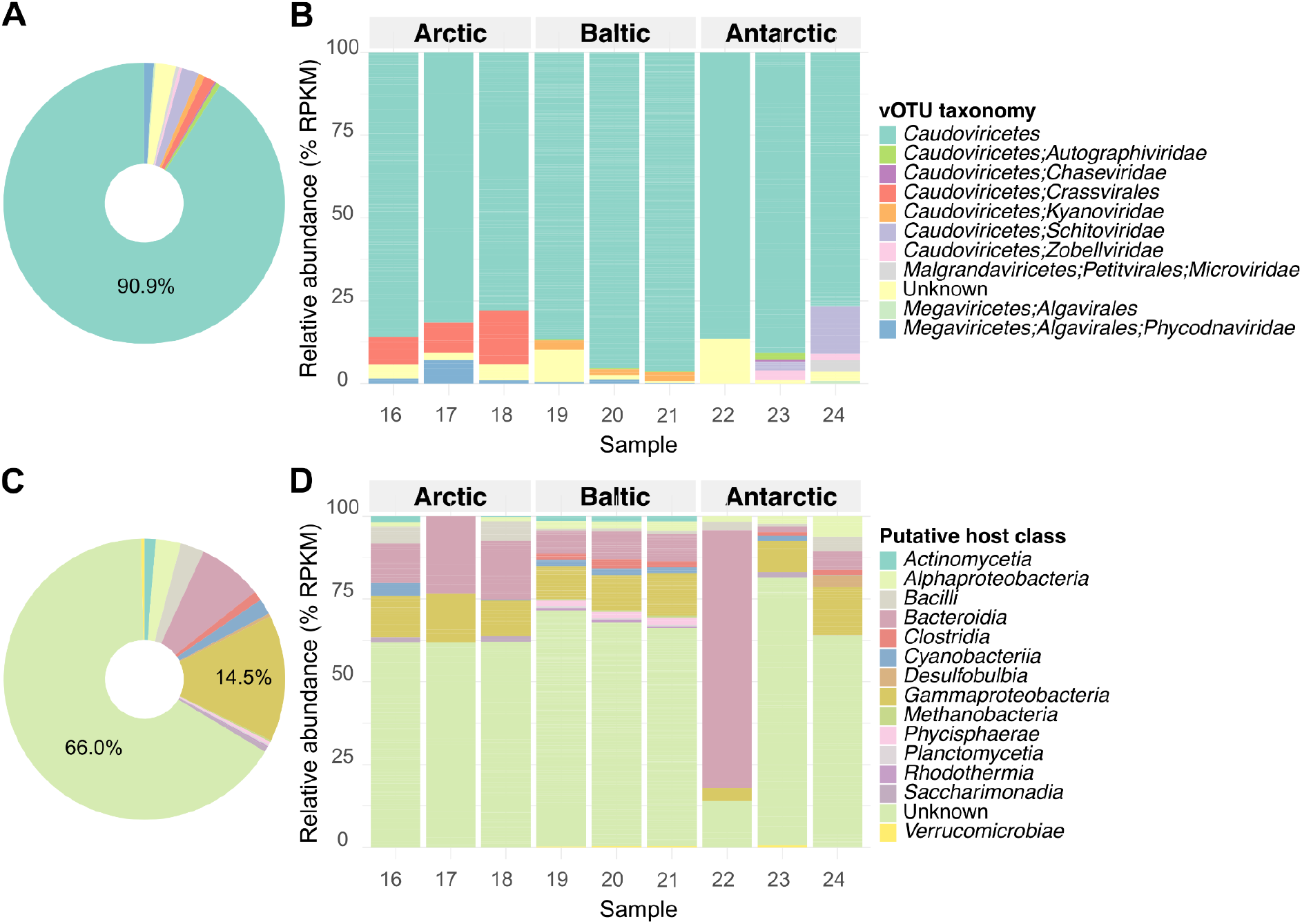
Predicted (A, B) viral taxa and (C, D) hosts for the obtained vOTUs. (A) and (C) show total counts for all vOTUs, while (B) and (D) show relative abundances of each viral taxon across samples. Color codes are the same for the upper and lower panels, respectively. In A and B, vOTUs taxonomy groups reflect the class- and/or lower-level predictions. In B and D, the horizontal coverage threshold of 0.1 applied. RPKM, reads per kilobase million.

The class *Malgrandaviricetes* included two vOTUs assigned to the family *Microviridae*, comprising tailless single-stranded (ss) DNA bacteriophages. Among the class *Megaviricetes*, the vOTUs were suggested to belong to the order *Algavirales* or further to the family *Phycodnaviridae*, i.e., representing eukaryotic (algal) viruses.

In terms of relevant abundance, vOTUs classified as *Caudoviricetes* were dominant across all samples, which was expected, since *Caudoviricetes* included the vast majority of all identified vOTUs (Fig. 3B). *Caudoviricetes* vOTUs that could be further classified to the order *Crassvirales* were abundant in samples 16 and 18 (Arctic). *Kyanoviridae* vOTUs were abundant in samples 19-21 (Baltic Sea), while *Schitoviridae* and *Zobellviridae* ones were observed in samples 23 and 24 (Antarctica). The taxonomic predictions presented here are generally in line with other reports, as the studied Arctic and Antarctic sea ice vOTUs could be classified mostly as *Caudoviricetes* (Zhong et al. 2020; Liu et al. 2022). In addition, *Caudoviricetes* have dominated in the Baltic Sea waters and sediments (Heyerhoff et al. 2022) and surface waters of Prydz Bay, Antarctica (Gong et al. 2018). A recent survey of DNA viruses in the Southern Ocean revealed extensive novel viral diversity, particularly highlighting previously unknown and diverse *Crassvirales* sequences (Piedade et al. 2024).

*Malgrandaviricetes, Megaviricetes*, and unclassified vOTUs were found only at low relative abundance, which reflects a low number of vOTUs assigned to these groups. *Microviridae* vOTUs were present only in the Antarctic sample 24. Microviruses are ubiquitous phages across environments (Kirchberger and Ochman 2023) and have dominated among ssDNA viruses in water samples from Prydz Bay in Antarctica (Gong et al. 2018). *Nucleocytoviricota* viruses have been found abundant in the Southern Ocean waters (Piedade et al. 2024), Baltic Sea surface waters (Heyerhoff et al. 2022), and the cold Arctic waters (Gao et al. 2021), but only at low abundance in Antarctic sea ice (Liu et al. 2022). Unlike in the previous study of the Antarctic sea ice, where filamentous phages (*Inoviridae*) were abundant in several samples (Liu et al. 2022), we observed no vOTUs classified as *Inoviridae*. Also, no virophages were found in our samples.

### Putative hosts of the identified vOTUs were mostly *Gammaproteobacteria* or *Bacteroidia*

Using iPHoP, 187 vOTUs were assigned to putative hosts: 186 to bacteria and only one to an archaeal host (Table S5). Thus, putative hosts for the majority of vOTUs (363) remained unknown. At the phylum level, most predictions (96) were for *Pseudomonadota*, followed by *Bacteroidota* (42), *Bacillota* (15), and *Cyanobacteriota* (10). Within *Pseudomonadota*, 80 and 16 predicted hosts belonged to the classes *Gammaproteobacteria* and *Alphaproteobacteria*, respectively (Fig. 3C). In terms of relative abundance, most samples were dominated by vOTUs with unknown hosts (Fig. 3D). Among vOTUs with predicted hosts, the ones assigned to the class *Gammaproteobacteria* and *Bacteroidia* were characterized by high relative abundance across samples. However, a noticeably high relative abundance of vOTUs linked to *Bacteroidia* in sample 22 is likely explained by a low number of vOTUs detected in the sample (see below). vOTUs linked to the class *Phycisphaerae* (phylum *Planctomycetota*) were observed only in the Baltic Sea ice samples.

The host phyla predicted for the most abundant vOTUs were among those with high relative abundance in the bacterial taxonomic profiling (Fig. 2A). In general, the host prediction results reinforce observations from other sea ice studies: among vOTUs *in silico* linked to their putative hosts, those associated with *Gammaproteobacteria* dominate in the Arctic and Antarctic sea ice (Liu et al. 2022; Zhong et al. 2020). Phages infecting *Gammaproteobacteria* have been abundant or even dominant also in the Southern Ocean waters (Piedade et al. 2024). In the Nordic Seas the most abundant viral populations among bacterial viruses were linked to *Pseudomonadota* hosts (Gao et al. 2021). The same study also showed that most virus-host linkages were positively related, highlighting covariations between viral and host communities in subarctic seas (Gao et al. 2021). vOTUs linked to *Bacteroidetes* (*Bacteroidota*) have also been shown abundant in the Arctic lower sea ice layers and sea ice brines (Zhong et al. 2020), and vOTUs linked to *Flavobacteriia* (*Bacteroidota*) have been abundant in Antarctic sea ice (Liu et al. 2022).

Using BACPHLIP, *in silico* lifestyle predictions suggested 118 and five vOTUs represented virulent and temperate phages, respectively, in our dataset (Table S5). In Phold-based annotations (see below), 106 and 23 vOTUs were predicted to encode genes related to lysis and/or integration/excision, respectively, including nine vOTUs with both types of genes predicted. Given that the lifestyle of the vast majority of vOTUs remained unknown and most vOTUs were incomplete, it is difficult to make conclusions of prevailing infection strategies across the samples.

Very little is known about viral infection strategies and virus-host dynamics in sea ice. *In silico* analyses suggest that up to ∼25% of viruses in Antarctic sea ice may adopt a lysogenic infection strategy, compared to at most ∼4% in the under-ice seawater (Cao et al. 2020). Notably, virus-host interactions are not restricted to lytic/lysogenic infections, but may also confer benefits to hosts (Roossinck 2011). For example, the filamentous sea ice phage f327 does not kill its host, *Pseudoalteromonas* spp., through cell lysis but instead decreases its growth rate and tolerance to NaCl and H_2_O_2_ while enhancing its motility and chemotaxis, which can benefit host survival in winter conditions and prevent over-blooming in summer (Yu et al. 2014). In addition, viruses can modulate their host metabolism through AMGs, with perhaps the best-known example being marine cyanophages that support their host’s photosynthesis by expressing genes that encode photosystem proteins (Lindell et al. 2004). Annotating sea ice vOTUs recovered in this study also revealed putative AMGs, including those involved in photosynthesis (see below).

### Sea ice vOTUs showed diverse ORFs including putative AMGs

geNomad-based gene annotation for seven vOTUs putatively classified as *Megaviricetes* included 70 ORFs, most of which (56) had unknown functions (Table S6). Annotating 543 non-eukaryotic vOTUs with Phold resulted in 6,613 predicted ORFs, averaging 1.3 ORFs per kilobase (Fig. 4A, Table S7). From all ORFs, 44% (2912) had unknown functions, while the rest were grouped into the following PHROG categories: head and packaging (925 ORFs), tail (816), DNA, RNA and nucleotide metabolism (791), connector (176), lysis (139), transcription regulation (103), and integration and excision (30). In addition, 482 ORFs were of other functions and 237 ORFs were annotated as moron, AMGs and host takeover, which we refer to as potential “AMGs” below. ORFs belonging to the latter category could be found in 138 vOTUs and mostly as 1-2 ORFs per vOTU, while 19 vOTUs contained 3-13 such ORFs.

**Figure 4.**
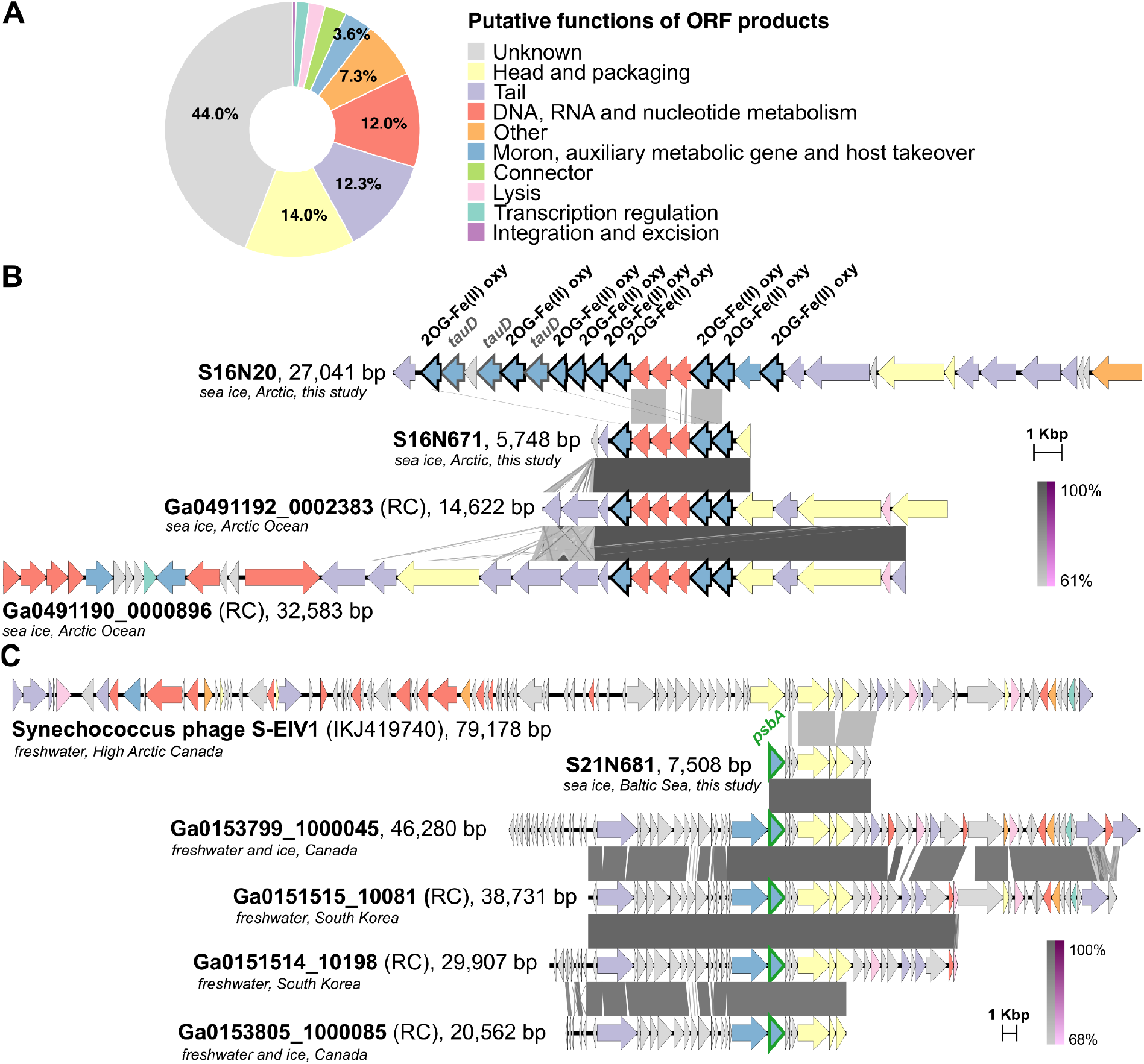
Annotated genes in vOTUs. (A) Functional categories predicted for putative ORF products across all 550 vOTUs. (B) The vOTUs S16N20 and S16N671 (this study), putatively encoding multiple 2OG-Fe(II) oxygenase genes (outlined in bold black) and taurine catabolism dioxygenase genes (*tauD*, outlined in bold grey), as well two related UViGs found in the IMG/VR database. (C) The vOTU S21N681 (this study), Synechococcus phage S-EIV1 (acc. no. KJ410740), and three related IMG/VR UViGs. Predicted *psbA* genes are outlined in bold green. All genomes were (re-)annotated with Phold. UViGs are labelled with their respective scaffold IDs. RC, reverse complement. Identities (BLASTn) between the genomes are shown as shades of grey (purple for inverted). The color key for ORF functions is the same for both sections.

The highest number of putative AMGs (13) was found in the vOTU S16N20, which was ∼27 kbp long, assembled from sample 16 (Arctic), and predicted as a tailed dsDNA virus belonging to the class *Caudoviricetes* with an unknown host. Notably, S16N20 contained nine putative 2-oxoglutarate-Fe(II) oxygenase (2OG-Fe(II) oxygenase) and three taurine catabolism dioxygenase (*tauD*) genes (Fig. 4B). These multiple genes were not identical, and the contig coverage suggested no apparent assembly artifacts. Another vOTU, S16N671, which was ∼6 kbp long, assembled from the same Arctic sample, also predicted as *Caudoviricetes* with no known host, had a set of three putative 2OG-Fe(II) oxygenase genes, similar to those found in S16N20. In addition, the search against the IMG/VR v.4 database retrieved two hits to UViGs that had a region that was 99% identical to S16N671, including three 2OG-Fe(II) oxygenase genes. According to the IMG/VR record, these UViGs were both *Caudoviricetes* with no predicted hosts and originated from melted sea ice microbial communities from the Arctic Ocean. The two vOTUs, S16N20 and S16N671, were grouped into the same genus cluster (see below), which contained no other vOTUs. Putative 2OG-Fe(II) oxygenase genes were predicted also in 19 other vOTUs from all three sampling regions.

2OG-Fe(II) oxygenase superfamily proteins catalyse various oxidative reactions involved in important cellular metabolic processes (van Staalduinen and Jia 2015). 2OG-Fe(II) oxygenase-encoding genes can be found in viral genomes, e.g., in viruses that infect green microalgae (Derelle et al. 2015) and in cyanophages (Sullivan et al. 2010; Brum et al. 2013; Ma et al. 2014). These genes can occur in multiple copies: for example 24 copies were identified in the genome of S-SCSM1, a T4-like cyanophage isolate that infects *Synechococcus* and *Prochlorococcus* strains (Wang et al. 2022). 2OG-Fe(II) oxygenases are suggested to play a role in phage DNA repair under nitrogen-limited conditions during cyanophage infection (Weigele et al. 2007; Sullivan et al. 2010). Furthermore, taurine dioxygenase, a member of the larger 2OG-Fe(II) oxygenase superfamily (Martinez and Hausinger 2015), is important for the utilization of taurine as a sulfur source under starvation conditions in bacteria (van der Ploeg et al. 1996). *tauD* genes are among abundant viral AMGs involved in organosulfur metabolism across various environments (Kieft et al. 2021) and were also found in *Caudoviricetes* sequences in the epishelf lake in Milne Fiord, the High Arctic (Labbé et al. 2022).

AMGs related to photosynthesis such as *psbA*, which encodes photosystem II D1 protein, and ferredoxin genes were identified in one and 16 vOTUs, respectively. The *psbA* gene product is a key component of oxygenic photosynthesis and is found in cyanophages (Sullivan et al. 2006; Zheng et al. 2013). The vOTU carrying *psbA* (S21N681) was 7.5 kbp long, assembled from sample 21 (Baltic Sea), unclassified, and had no host links. However, when the putative *psbA* gene sequence was searched against the NCBI core_nt database, multiple hits to cyanophages were retrieved. Using the whole S21N681 sequence as the query, one of the best matches was to the complete genome of Synechococcus phage S-EIV1 (acc. no. KJ410740, coverage: 76%, identity: 75%). Notably, this match did not include the *psbA* gene, which is apparently absent in S-EIV1 (Fig. 4C). S-EIV1 was isolated from freshwaters in High Arctic Canada on the polar *Synechococcus* sp. PCCC-A2c and represents a group of related cyanophages that are widespread in aquatic environments (Chénard et al. 2015). Indeed, a search against the IMG/VR database retrieved many hits (90 hits of >50% query coverage and >77% identity) to viral sequences in geographically distant aquatic environments, but the best hits (>75% query coverage and > 96% identity) were to vOTUs found in microbial communities from Lake Erie water and ice samples (Ontario, Canada), Thompson River surface water (Kamloops, Canada), Lake Soyang (South Korea), and Han River (South Korea) (all shown in Fig. 4C), which are classified as *Caudoviricetes* with no known hosts. From 16 vOTUs, where putative ferredoxin-encoding genes were predicted, two vOTUs were linked to *Cyanobacteriota* hosts. Ferredoxin genes are also widespread in cyanophages and support electron transfer to host-encoded oxidoreductases involved in sulfur assimilation (Campbell et al. 2020).

Among other putative AMGs, those involved in queuosine biosynthesis (e.g., *queC*-, *queD*-, and *queE*-like), cobalamine biosynthesis, cysteine metabolism (*cysH*), and biosynthesis of lipopolysaccharides belong to the apparently globally conserved pool of viral genes found across different environments (Kieft et al. 2020). In addition, gene products that may regulate transcription and translation in the host to support cell survival under stress conditions, such as MazG-like pyrophosphatase, PhoH-like phosphate starvation-inducible, heat shock, and cold shock proteins, were identified. Various putative proteins involved in virus-host arm races were also found: DNA methyltransferases (66 predictions), anti-restriction proteins (e.g., DarB-like), as well as anti-CRISPR proteins, and several bacterial virulence factor proteins, including in vOTUs putatively assigned to *Enterobacteriaceae* and *Flavobacteriaceae* hosts. Overall, a wide variety of virus-host interactions could be inferred from the predicted functional potential in sea ice vOTUs.

### Viral communities differed across samples, but shared some genus-or family-level similarities within this and other reference datasets

In the PCoA analyses of viral communities, the Baltic Sea samples clustered tightly together, being distant from the rest of the samples (Fig. 5A). However, more samples would be needed to verify this observation. Samples 17 and 22, which were flooded, had the lowest number of vOTUs present: 23 and 9 vOTUs were detected in samples 17 and 22, respectively. In addition to the possible impact of flooding, sample 22 had the lowest number of reads among the studied samples, and since virus sequences usually constitute only a small portion of all sequences in bulk metagenomes (Roux et al. 2021), the viral DNA in this sample might have stayed undersequenced.

**Figure 5.**
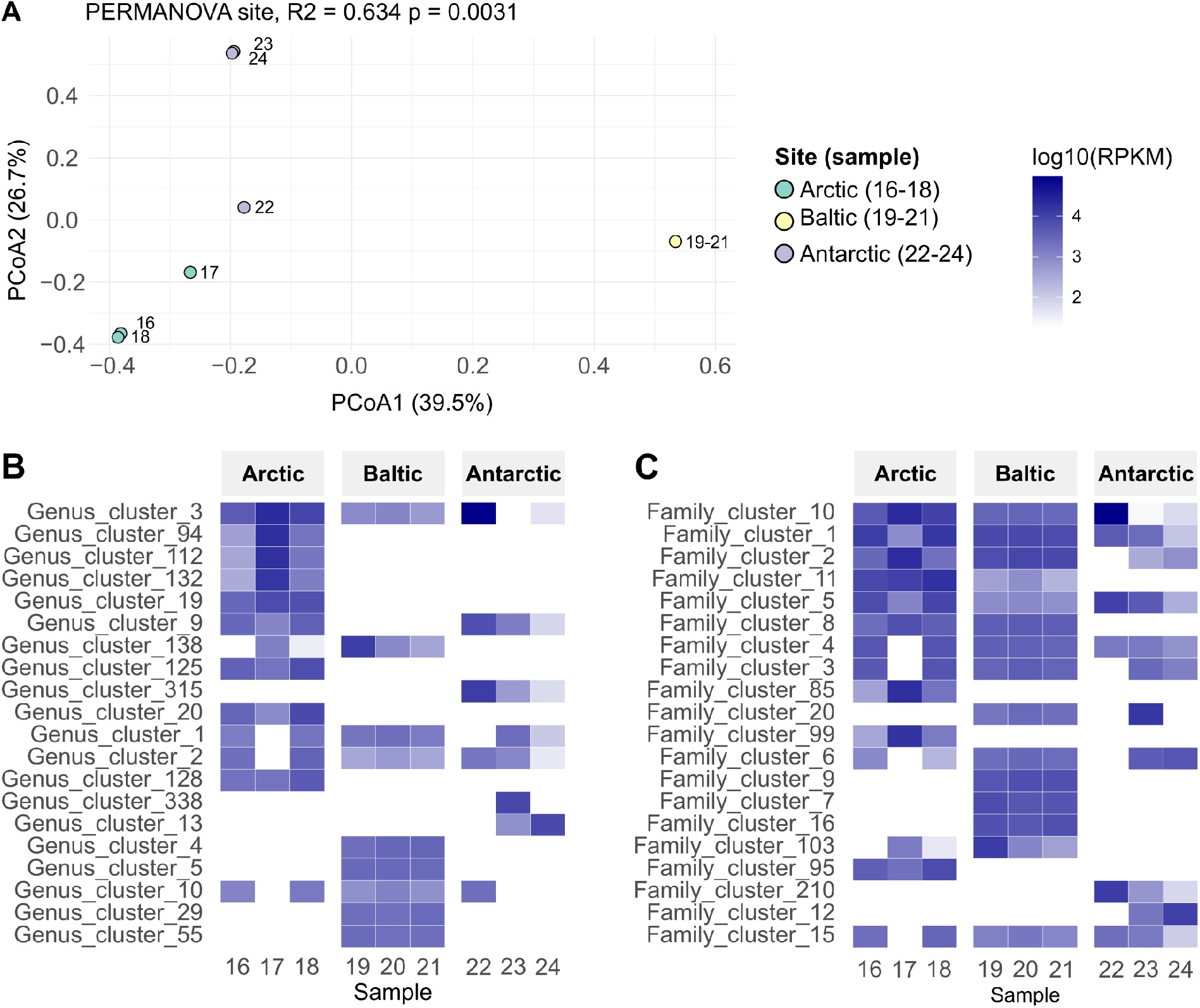
Viral community composition. (A) Principal coordinates analysis (PCoA) of the vOTUs, samples are color-coded (see inset). Percentages in brackets indicate the percentage of the total variation in the data that is explained by that axis. Heatmaps showing the coverage of the 20 most abundant (B) genus and (C) family clusters across samples. The horizontal coverage threshold of 0.1 applied for all analyses here.

Individual vOTUs were shared mainly only within their sampling regions (Fig. S3). However, when vOTUs were clustered into 405 genus- and 275 family-level groups (Table S5), some clusters could be observed across three sampling regions (Fig. 5BC). For example, genus cluster 3 (comprising seven vOTUs and being part of family cluster 10) was present in all samples, except sample 23 (Fig. 5B), and family cluster 1 (comprising 37 vOTUs) was present across all nine samples (Fig. 5C). The largest clusters comprised *Caudoviricetes* and/or unclassified vOTUs. The two detected microvirus vOTUs were grouped into the same genus, while seven *Megaviricetes* vOTUs belonged to six different genus clusters (Table S5).

To further explore genus-or higher-level similarities between vOTUs within our dataset and other references, we performed a whole-genome gene-sharing network analysis with VConTACT2, using vOTUs from this study, ice-associated UViGs retrieved from the IMG/VR database (Tables S8, S9), vOTUs from Arctic sea ice and cryopeg brines (Zhong et al. 2020), vOTUs from Antarctic under-ice water (Lopez-Simon et al. 2023), as well as sequences from NCBI GenBank including the sea ice virus isolates (Table S9, Fig. 6). From the 141 vOTUs obtained in this study (>10 kbp), more than half (77) stayed unassigned to any cluster, while 64 vOTUs clustered with each other and/or other references. Overall, the vOTUs from this study clustered more within their own sampling region rather than the other two: from 18 viral clusters (VCs, genus-level groupings) that contained two or more vOTUs from this study, 12 had vOTUs from only the same sampling region, while the other six VCs included vOTUs from two regions. These inter-region clusters had either Arctic and Antarctic or the Arctic and Baltic vOTUs clustered, but not Antarctic with Baltic ones together. Nonetheless, higher-level (family-level or higher) links between vOTUs from all three sampling regions could still be observed as connections in the network (e.g., see the dash-line square in Fig. 6).

**Figure 6.**
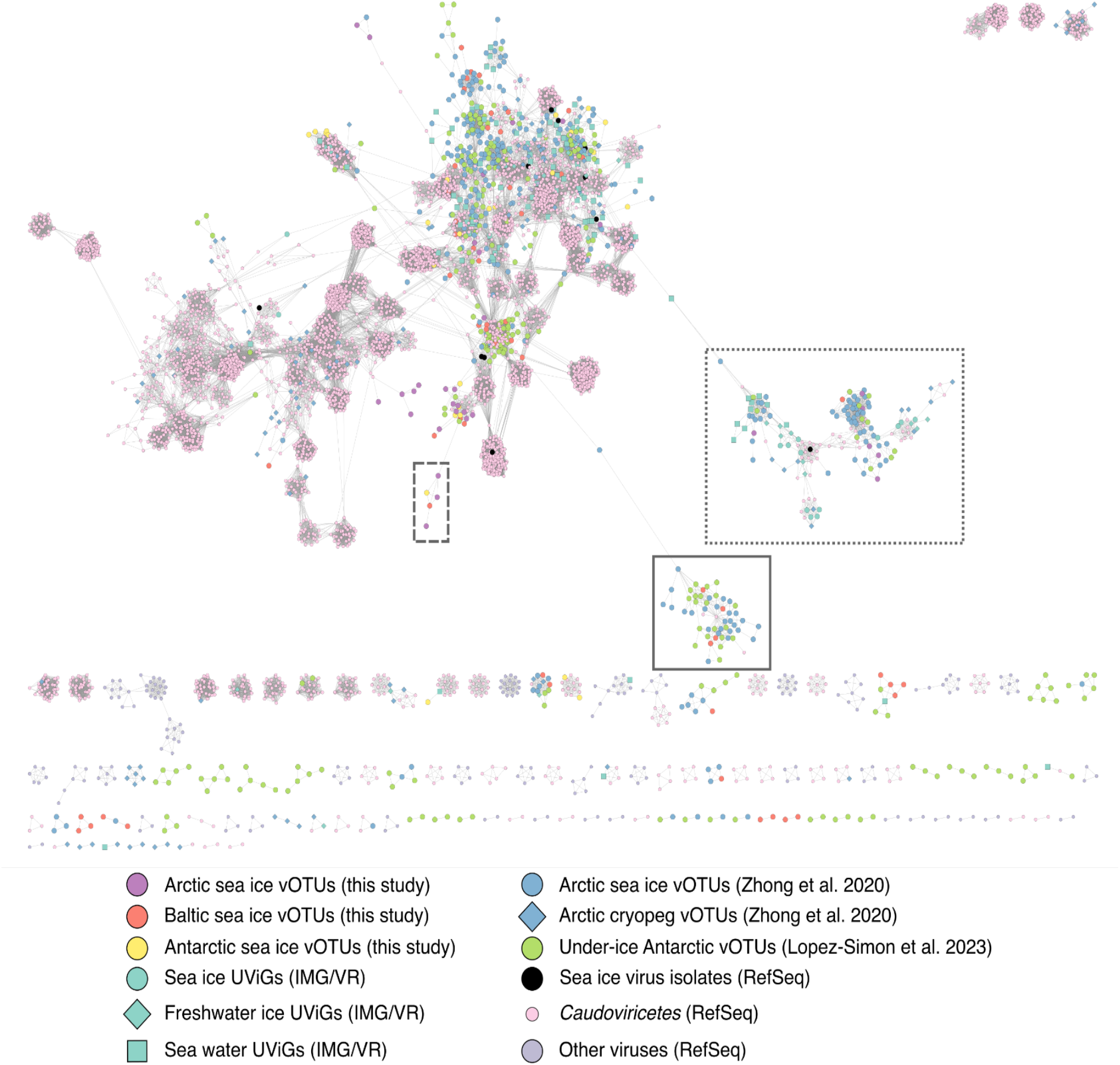
VConTACT2 network of the viral clusters. Viral genomes and vOTUs are shown as color-coded circles, squares, or diamonds. The dash-line box outlines an example of the connections between this study’s vOTUs from all three sampling regions; the solid-line box shows an example of the connections observed between this study and other reference datasets; and the dotted-line box depicts the connections between Flavobacterium phage 11b (Arctic sea ice isolate, GenBank acc. no. AJ842011.2) and other vOTUs/UViGs.

Ten VCs contained vOTUs exclusively from this study, while 19 VCs included vOTUs from this study and Arctic vOTUs from (Zhong et al. 2020), and 14 VCs had vOTUs from this study and Antarctic vOTUs from (Lopez-Simon et al. 2023). vOTUs from this study clustered primarily with Arctic sea ice vOTUs rather than cryopeg ones reported in (Zhong et al. 2020), as only three VCs contained cryopeg vOTUs, of which two VCs also contained sea ice vOTUs from the same study (Zhong et al. 2020). Other studies have suggested that viral communities in Arctic sea ice differ from those in Arctic permafrost cryopegs (Zhong et al. 2020, 2023), but share similarities with the underlying seawater (Zhong et al. 2023).

VConTACT2 clustering was observed across geographic distances, as Arctic/Baltic vOTUs from this study shared some clusters with Antarctic under-ice vOTUs from (Lopez-Simon et al. 2023), and this study Antarctic vOTUs also shared some clusters with Arctic sea ice vOTUs from (Zhong et al. 2020). The solid-line square in Fig. 6 shows an example of genus-or higher-level connections between Baltic vOTUs from this study, Arctic vOTUs from (Zhong et al. 2020) and Antarctic under-ice vOTUs from (Lopez-Simon et al. 2023).

Eight VCs included both vOTUs from this study and ice-associated UViGs from the IMG/VR database. These UViGs were from melted sea ice in the Arctic Ocean, sea ice brine near Alaska, or Tibetan glacier (see Table S8 for the detailed information on UViGs and Table S9 for their clusters). In addition, vOTUs from this study shared six VCs with RefSeq sequences, including two VCs with sea ice virus isolates (see below for isolates) and one VC containing the vOTU S21N312 and the cyanophage Synechococcus phage S-CBWM1 (Xu et al. 2018).

Other studies have shown that polar sea ice viruses tend to share most similarities within their regions. In (Zhong et al. 2023), Arctic sea ice vOTUs clustered with other Arctic viruses from the global oceans (GOV2) dataset. In (Liu et al. 2022), Antarctic sea ice vOTUs were mostly endemic, with only a fraction of sequences matching those in other ecosystems (Liu et al. 2022). Our observations suggest that some limited genus-or higher-level genetic similarities between (sea) ice vOTUs could still be found across geographic distances.

### Limited similarities were observed between vOTUs and known sea ice virus isolates

Mapping metagenomic reads to the complete genomes of 11 sea ice virus isolates (Table S2) resulted in very low alignment rates, close to 0%. Nonetheless, when the sea ice virus isolate genomes were compared to the metagenomic contigs obtained here using BLASTn or BLASTp, relatively short regions of similarities were found between the Antarctic contigs and Antarctic sea ice phage isolates PANV2 and OANV2 (Luhtanen et al. 2018; Demina et al. 2022) (Fig. S4). Similarities were also observed between the contigs from the Arctic sample 16 and the Flavobacterium phage 11b, which was originally isolated from Arctic sea ice (Borriss et al. 2003). However, not all of these contigs (Fig. S4) were predicted by geNomad as viral.

In the gene-sharing network analysis by vConTACT2 (Fig. 6), the Antarctic phage isolates PANV2 and OANV1 were the only ones that clustered with the vOTUs assembled from this study (Antarctic samples). Both PANV2 and OANV1 also clustered with some Arctic melted sea ice UViGs in their respective VCs. In addition, OANV1 clustered with RefSeq sequences (phages of *Bordetella, Enterobacter, Escherichia, Pseudomonas, Salmonella*, and *Xanthomonas*), under-ice Antarctic vOTUs (Lopez-Simon et al. 2023), and Arctic vOTUs (Zhong et al. 2020). Although the rest of sea ice isolates shared no VCs with the vOTUs from this study, we observed their connections to other reference datasets. The Antarctic isolate OANV2 clustered with one Arctic IMG/VR UViG and one Arctic sea ice vOTU (Zhong et al. 2020). Among Baltic Sea ice virus isolates, Shewanella sp. phages 1/4 and 1/40 clustered together, consistent with the suggestion that they belong to the same genus (Senčilo et al. 2015). Shewanella sp. phage 1/41 shared a VC with one Arctic IMG/VR UViG and one Arctic sea ice vOTU (Zhong et al. 2020). Flavobacterium sp. phage 1/32 clustered with two other *Flavobacteriaceae* phages and one Arctic cryoconite UViG. Shewanella sp. phage 1/44 formed a VC with Stenotrophomonas phage S1 and an Arctic cryopeg vOTU (Zhong et al. 2020). The Arctic sea ice virus isolate Flavobacterium phage 11b was categorized as an outlier, but in the network, it is located next to other *Flavobacterium* phage sequences. The dotted-line square in Fig. 6 shows a group of genus-or higher-level connections between Flavobacterium phage 11b, *Caudoviricetes* sequences (RefSeq) for *Bacteroides, Cellulophaga, Flavobacterium, Nonlabens, Polaribacter*, and *Riemerella* phages, as well as Arctic and Baltic vOTUs from this study and other Arctic (Zhong et al. 2020) and Antarctic (Lopez-Simon et al. 2023) vOTUs.

Previously, the Arctic isolate Flavobacterium phage 11b was found abundant in surface waters in Prydz Bay, Antarctica (Gong et al. 2018). Viral sequences related to the Antarctic phages OANV1 and PANV2 were also observed in geographically distant aquatic environments (Demina et al. 2022). Furthermore, reads mapping to the structural proteins in the Baltic Sea ice virus isolates could be found from other environments (Senčilo et al. 2015). Thus, some sea ice virus isolates may represent more widespread viral groups found in various distant environments.

## Conclusions

Metagenomic analyses of nine sea ice samples from the Arctic, Antarctic, and Baltic Sea revealed diverse unicellular organisms and viruses across samples. A total of 550 vOTUs were recovered in bulk sea ice metagenomes. The majority of vOTUs were putatively classified within the class *Caudoviricetes*, which is consistent with other reports on the diversity of viruses in sea ice and polar environments (Zhong et al. 2020; Liu et al. 2022; Gong et al. 2018; Piedade et al. 2024; Kanaan and Deming 2025). About 34% of the vOTUs were *in silico* linked to their putative bacterial hosts, primarily from the classes *Gammaproteobacteria* and *Bacteroidia*, which are among dominant bacterial taxa previously observed in such environments (Rapp et al. 2025). Still, the samples were dominated by viruses with unknown hosts. Nonetheless, predicted virus-host linkages suggest that viruses play an important role in shaping their host communities in sea ice, as they are associated with abundant sea ice bacteria.

The viral genomes encoded products with diverse putative functions, including potential AMGs involved in oxidative metabolism (multiple 2OG-Fe(II) oxygenase genes), photosynthesis (*psbA* and ferredoxin genes), and stress response (*tauD* genes and others). Using the IMG/VR database, related viral genomes carrying similar 2OG-Fe(II) oxygenase or *psbA* genes could be found in freshwater, marine and ice environments from the Arctic and more distant geographical locations. Other predicted functions, such as those related to defense mechanisms, point to diverse and complex virus-host interactions in sea ice. Notably, almost half of the predicted genes had unknown functions, highlighting that the genetic diversity of sea ice viruses remains largely unexplored.

While no vOTUs were shared across all nine samples, family-level similarities were observed throughout. Comparison with reference datasets (Zhong et al. 2020; Lopez-Simon et al. 2023) and high-quality ice-associated UViGs retrieved from IMG/VR revealed connections to other marine and ice viral communities, while similarities to cultured sea ice virus isolates remained very limited. This study contributes to expanding the sequence space of sea ice viruses, including those in Antarctica, a logistically challenging region. Still, more sampling efforts are needed to resolve the geographic diversity and ecological roles of sea ice viruses in polar and subpolar environments.

## Funding

This work was supported by the Research Council of Finland (330977 to T.D., 354462 to J.H., 276739 to H.K.); the Research Council of Norway (244646 to P.A.); and the Kone Foundation (to T.D.). P.A. also acknowledges the Centre for ice, Cryosphere, Carbon and Climate (iC3) supported by the Research Council of Norway through its Centres of Excellence funding scheme (332635). The work conducted by the U.S. Department of Energy Joint Genome Institute (https://ror.org/04xm1d337), a DOE Office of Science User Facility, is supported by the Office of Science of the U.S. Department of Energy operated under Contract No. DE-AC02-05CH11231. H.M.O. was supported by the University of Helsinki and the Research Council of Finland by funding for FINStruct and Instruct Centre FI, part of Biocenter Finland and Instruct-ERIC and by Horizon MSCA 101120407.

## Acknowledgements

We thank CSC Finland for computational resources and technical support. We acknowledge the DNA Sequencing and Genomics Laboratory, Institute of Biotechnology, University of Helsinki, for sequencing. Anthropic Claude Sonnet 4 was used for the development of R codes and Grammarly was used to enhance grammar and clarity of the manuscript text. All outputs generated by these tools were reviewed and verified by the authors.

## Conflict of interests

The authors declare no conflict of interests.

## Data availability

The reads are available from ENA: ebi.ac.uk/ena/browser/view/PRJEB101482. vOTU and TE sequences identified in this study can be downloaded from Figshare: https://doi.org/10.6084/m9.figshare.30437099.v1 for vOTUs and https://doi.org/10.6084/m9.figshare.30437210.v1 for other TEs.

## Notes

https://doi.org/10.6084/m9.figshare.30437099.v1

https://doi.org/10.6084/m9.figshare.30437210.v1

### Competing Interest Statement

The authors have declared no competing interest.

### Summary of Updates

Minor revisions in Materials & Methods and Results: - sampling and sample handling procedures clarified; - hypothesis and expected results clarified; - viral lifestyles in silico predicted; - AMG analyses clarified; - Phold predictions retained for non-eukaryotic viruses only (Figure 4 updated); - additional supplementary table for geNomad-based gene annotations for eukaryotic vOTUs added; - a few new references added.

