## Supplementary Figures for "Viral genetic diversity and functional potential in polar and subarctic sea ice"

[*]Shared first authorship

[1]Department of Microbiology, Faculty of Agriculture and Forestry, University of Helsinki, Helsinki, Finland

[2]Natural Resources Institute Finland (Luke), Helsinki, Finland

[3]Helsinki Institute of Sustainability Science (HELSUS), University of Helsinki, Helsinki, Finland

[4]DOE Joint Genome Institute, Lawrence Berkeley National Laboratory, Berkeley, CA, USA

[5]Finnish Environment Institute (Syke), Helsinki, Finland

[6]Department of Molecular and Integrative Biosciences, Faculty of Biological and Environmental Sciences, University of Helsinki, Helsinki, Finland

[7]Norwegian Polar Institute, Fram Centre, Tromsø, Norway

### Supplementary Figures

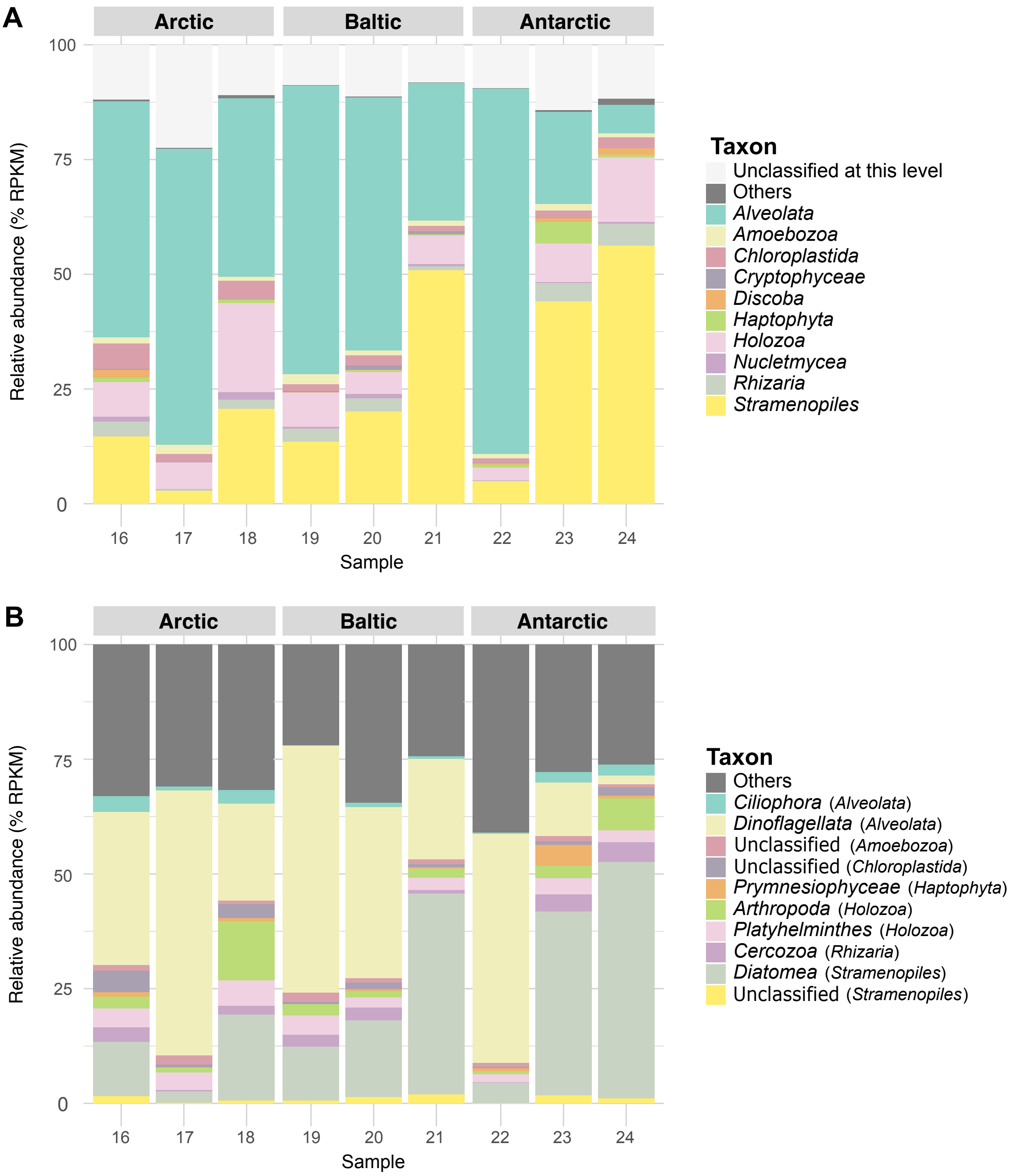


**Figure S1.** The ten most abundant eukaryotic taxa across samples, summarized at the (A) second and (B) third levels according to phyloFlash. The data were normalized after removing archaeal and bacterial sequences (see Materials and Methods).

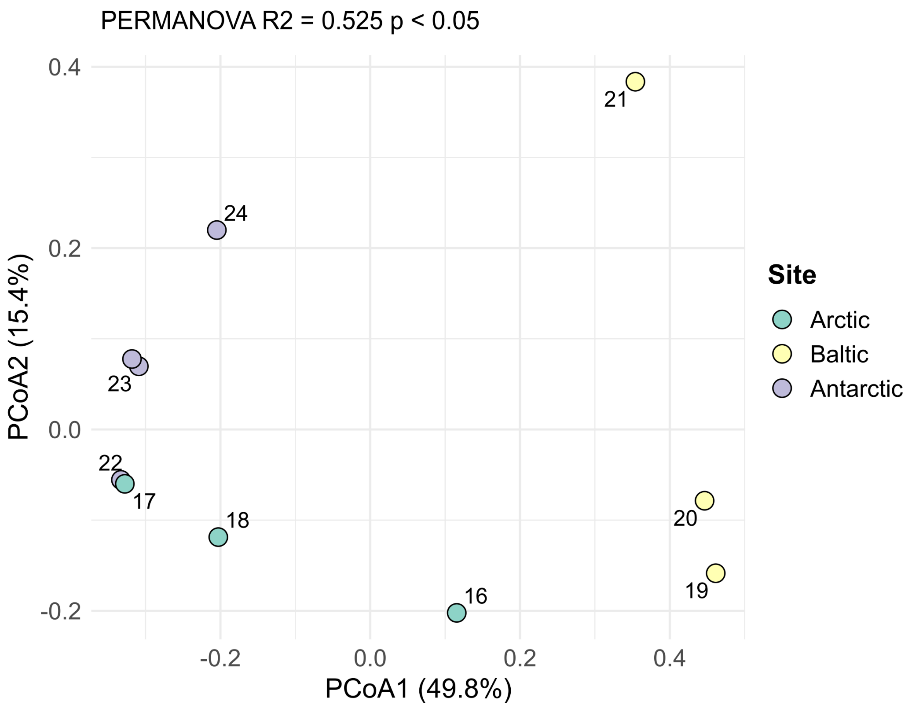


**Figure S2.** PCoA of the bacterial and archaeal community structure. The samples are colored based on their region. Percentages in brackets indicate the percentage of the total variation in the data that is explained by that axis.

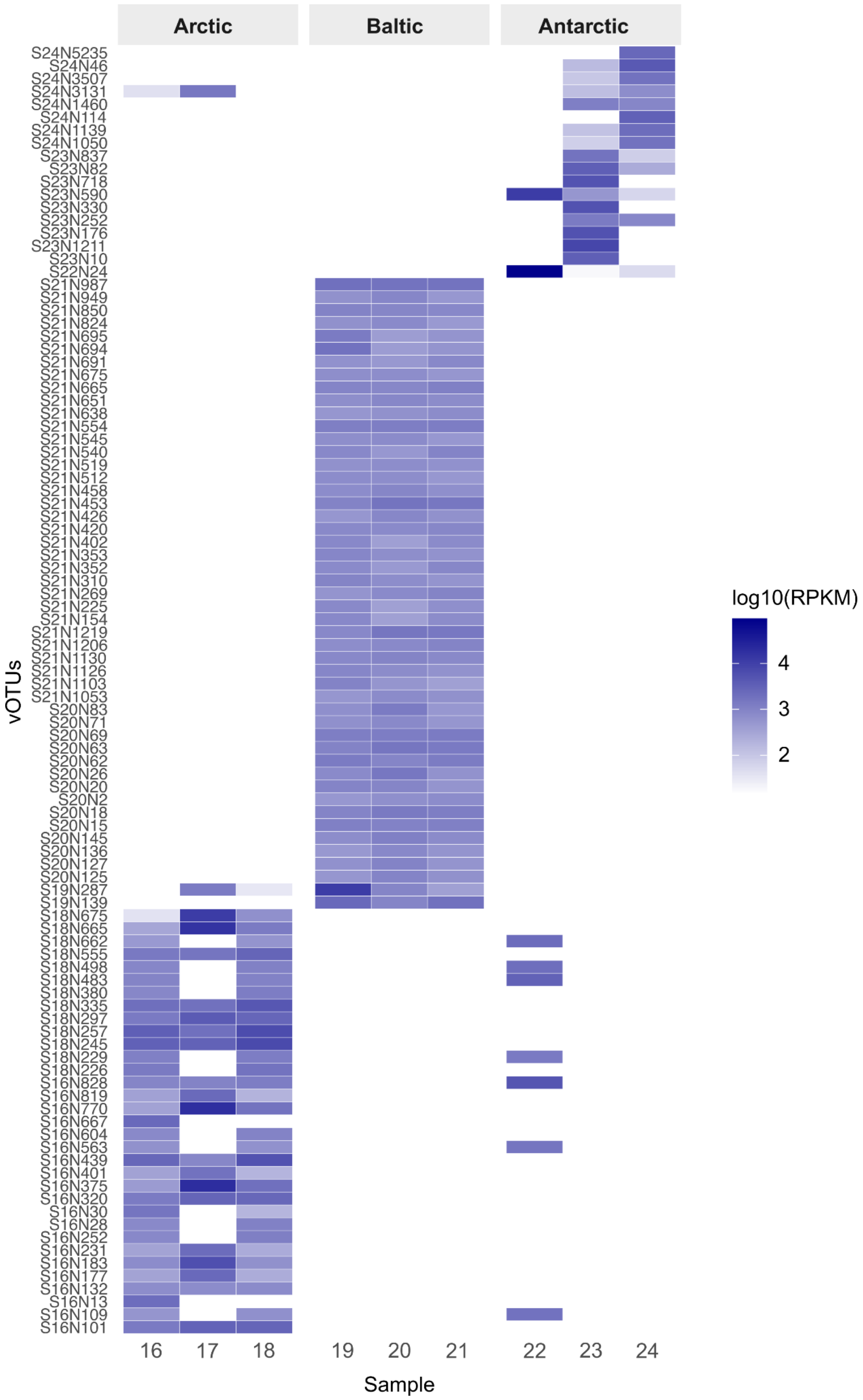


**Figure S3.** Heatmap showing the coverage of the 50 most abundant vOTUs across samples. The horizontal coverage threshold of 0.1 was applied.

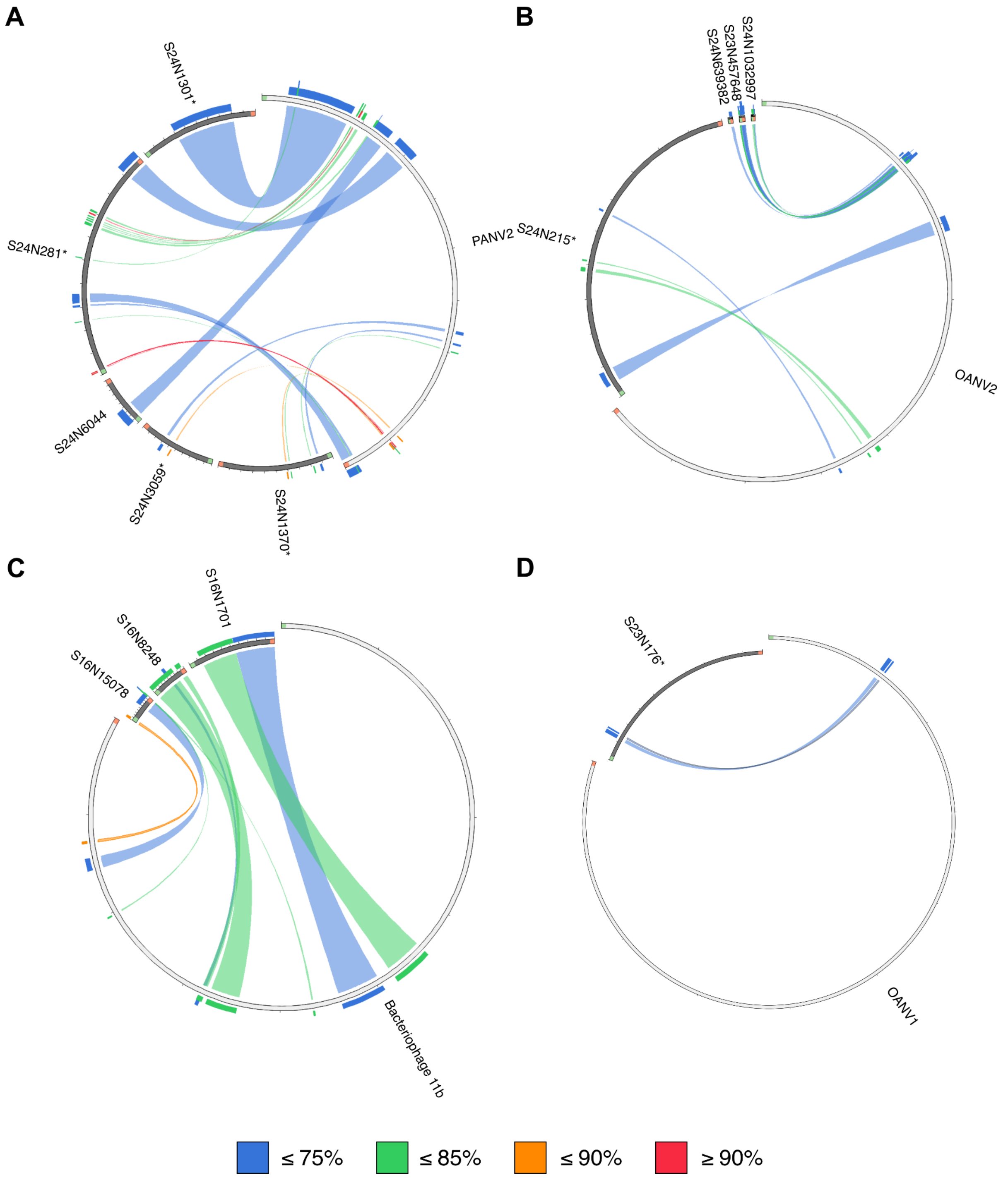


**Figure S4.** Similarities that the sea ice isolates share with the vOTUs and metagenomic contigs from this study. (A) PANV2 compared to Antarctic vOTUs and contigs, (B) OANV2 compared Antarctic vOTUs and contigs, (C) Flavobacterium phage 11b compared to Arctic contigs, and (D) OANV1 compared to one Antarctic vOTU. The names of vOTUs are followed by an asterisk (while metagenomic contigs that were not recognized as vOTUs lack it). The similarities are shown as % identity. The minimum and maximum % identity for each figure are: (A) 65.74%-93.75%, (B) 68.75%-78.72%, and (C) 72.69%-88.61%, and (D) 69.76%-72.41%. The start of a sequence is colored green and the end is red.
